# Polyamine Depletion by D, L-alpha-difluoromethylornithine Inhibits Ewing Sarcoma Metastasis by Inducing Ferroptosis

**DOI:** 10.1101/2024.06.14.599064

**Authors:** Rachel Offenbacher, Kyle W. Jackson, Masanori Hayashi, Jinghang Zhang, Da Peng, Yuqi Tan, Tracy Murray Stewart, Paul Ciero, Jackson Foley, Robert A. Casero, Patrick Cahan, David M. Loeb

## Abstract

Polyamine metabolism and signaling play important roles in multiple cancers but have not previously been studied in Ewing sarcoma. Here, we show that blocking polyamine synthesis with D, L-alpha-difluoromethylornithine (DFMO) causes a G1 cell cycle arrest, dose-dependent decreases in sarcosphere formation from Ewing sarcoma cell lines growing in non-adherent conditions and a decrease in clonogenic growth in soft agar. Further, we utilized our orthotopic implantation/amputation model of Ewing sarcoma metastasis to demonstrate that DFMO slowed primary tumor growth in addition to limiting metastasis. RNA sequencing demonstrated gene expression patterns consistent with induction of ferroptosis caused by polyamine depletion. Induction of ferroptosis was validated in vitro by demonstrating that ferrostatin-1, an inhibitor of ferroptosis, allows sphere formation even in the presence of DFMO. Collectively, these results reveal a novel mechanism by which DFMO prevents metastasis – induction of ferroptosis due to polyamine depletion. Our results provide preclinical justification to test the ability of DFMO to prevent metastatic recurrence in Ewing sarcoma patients at high risk for relapse.

## Introduction

Prior to the introduction of systemic chemotherapy for the treatment of localized Ewing sarcoma, event-free survival (EFS) rates were dismal (1). Systemic chemotherapy initially improved 5-year event-free survival rates to 60% (2), and with increasingly intense chemotherapy regimens the most recent Children’s Oncology Group trial demonstrated 78% EFS at 5 years (3). In contrast, metastatic disease at presentation has remained difficult to treat, with six-year survival rates of approximately 30% (4) that have not significantly changed in decades. Patients with recurrent disease face the most grim prognosis, with 5-year survival rates of only 10-15% (5), also unchanged despite numerous clinical trials, demonstrating the dire need for more effective therapies to prevent or treat relapsed and metastatic disease.

To gain a deeper understanding of the mechanisms underlying Ewing sarcoma metastasis, we developed an orthotopic implantation model utilizing patient-derived xenografts (PDX) (6). Intact fragments of Ewing sarcoma xenografts implanted within the pre-tibial microenvironment (orthotopic) spontaneously metastasize. Conversely, tumors implanted in the subcutaneous space do not metastasize, demonstrating that xenograft metastatic potential is critically dependent on the microenvironment. In exploring the biological underpinnings of this observation, we discovered that arginase, the enzyme responsible for metabolizing arginine to ornithine and toward polyamine production, is more strongly expressed in subcutaneous tumors than in those implanted orthotopically. This finding led us to further investigate the role polyamine signaling might play in Ewing sarcoma growth and metastasis.

Polyamines play a critical role in cell proliferation (7) and therefore have been investigated in a variety of different cancers. Colon (8), skin (9), prostate (10), and breast cancer (11) have all demonstrated increased polyamines, and polyamines have been shown to play a role in carcinogenesis (12), angiogenesis (13), tumor proliferation (14), and metastasis (15). The precise mechanism by which polyamines modulate these processes, however, remains unclear. Attempts at pharmacologic manipulation of this pathway have focused on targeted enzyme inhibition (16) and polyamine analogs (17). D, L-alpha-difluoromethylornithine (DFMO) is an irreversible inhibitor of ornithine decarboxylase (ODC)(18), the initial, rate-limiting enzyme within the polyamine biosynthetic pathway, and has been studied in a number of different cancers and investigated as a chemopreventive agent (19-22). DFMO administered to patients with high risk neuroblastoma after immunotherapy prolongs both event-free (EFS) and overall survival (OS) (23), and *in vitro* work has demonstrated a potential role for polyamines in neuroblastoma cancer stem cell differentiation (24). Informed by these results, we investigated the role of polyamines in Ewing sarcoma proliferation and metastasis and whether blocking the polyamine synthetic pathway with DFMO had therapeutic potential in the treatment of Ewing sarcoma. We found that inhibiting polyamine synthesis *in vitro* leads to a profound suppression of Ewing sarcoma cell clonogenicity and stem cell-like behavior. Additionally, utilizing our novel *in vivo* model of metastasis, we demonstrate that inhibiting polyamine synthesis limits Ewing sarcoma tumor growth and metastasis in a dose-dependent manner. Importantly, RNA sequencing demonstrated gene expression changes consistent with the model that polyamine depletion induces ferroptosis. Taken together, our findings demonstrate a novel insight into the mechanism of Ewing sarcoma metastasis, credential DFMO as a promising and readily translatable therapeutic treatment option for Ewing sarcoma patients at high risk of metastatic recurrence, and provide novel mechanistic insight into the role of polyamines in modulating ferroptosis.

## Results

### Quantification of polyamine levels in Ewing sarcoma tumors implanted orthotopically or subcutaneously

Our group has previously demonstrated that arginase expression varies in Ewing sarcoma PDXs depending on their implantation site and associated microenvironment, with higher levels in subcutaneous tumors compared with orthotopic tumors (6). To validate that the increase in arginase expression results in a downstream increase in polyamine synthesis, ODC activity and pooled polyamine levels were quantified in EWS4 patient-derived xenografts implanted either subcutaneously or in the orthotopic pretibial space. As anticipated, we observed significantly less ODC activity in orthotopically implanted tumors, 71.75 ± 8.6 pmol CO_2_/hr/mg protein, compared to those in the subcutaneous microenvironment, 137.9 ± 11.2 pmol CO_2_/hr/mg protein (p=0.0009) (Figure 1A). Further, we observed significantly less putrescine (PUT), spermidine (SPD), and spermine (SPM) in orthotopic tumors (Figure 1B), commensurate with the decreased ODC activity. Overall, these results confirmed that non-metastatic, subcutaneous tumors have significantly different polyamine metabolism compared to those implanted in the orthotopic microenvironment conducive to metastatic spread. Informed by this finding, we proceeded to investigate the effect of DFMO, an inhibitor of ODC, on Ewing sarcoma cells with a cancer stem cell phenotype, potentially targeting the subpopulation of cells thought to be responsible for metastasis and relapse.

**Figure 1:**
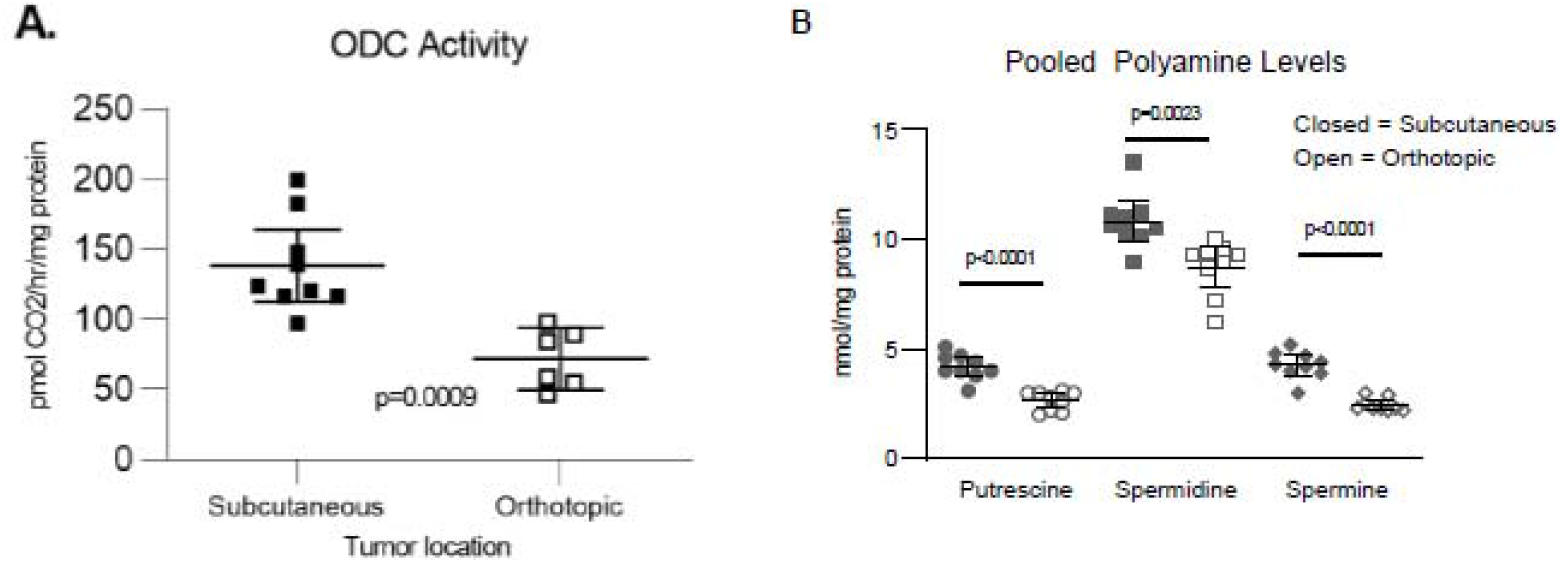
Ornithine decarboxylase activity and polyamine levels within orthotopic and subcutaneous Ewing sarcoma patient derived xenografts. (A) ODC activity was quantified in EWS4 patient-derived xenografts implanted subcutaneously or orthotopically. Error bars represent 95% confidence interval for triplicate evaluation of each tumor. Pooled data from 3 individual tumors, each analyzed in triplicate assays demonstrate a statistically significant difference (p=0.0009). (B) Levels of putrescine, spermidine and spermine were quantified in EWS4 patient-derived xenografts implanted orthotopically (open figures) and subcutaneously (closed figures). Error bars represent 95% confidence interval and p values were calculated using unpaired Student’s T-test.

### Blocking polyamine synthesis inhibits Ewing sarcoma proliferation and cancer stem cell phenotype in vitro

Our initial experiments investigating the effects of blocking polyamine synthesis with DFMO on Ewing sarcoma cell growth *in vitro* demonstrated dose-dependent inhibition of proliferation (Figure 2a-c). To determine whether the inhibition of proliferation reflects a cytostatic or cytotoxic effect, wash-out experiments were performed. Cells were cultured for 24 hours in regular medium followed by 4 days in DFMO. After 4 days, DFMO was removed and proliferation was evaluated in regular growth medium over the ensuing 6 days. Removal of DFMO allowed resumption of cell proliferation (Figure 2d-f), except at the highest doses, consistent with a cytostatic effect. Finally, to confirm that the effect of DFMO is mediated through depletion of intracellular polyamines, a spermidine add-back experiment was performed. Briefly, after culturing cells for 3 days in DFMO, spermidine was added to the cultures, along with aminoguanidine. The aminoguanidine is necessary to inhibit serum amine oxidase, present in fetal bovine serum, that would otherwise degrade spermidine and generate H_2_O_2_, aldehyde, and ammonia, all of which are cytotoxic (25). We found that adding back spermidine to cells treated with DFMO induces renewed proliferation despite the continued presence of drug (Figure 2g-i), consistent with DFMO-mediated polyamine depletion being the direct cause of proliferation arrest.

**Figure 2:**
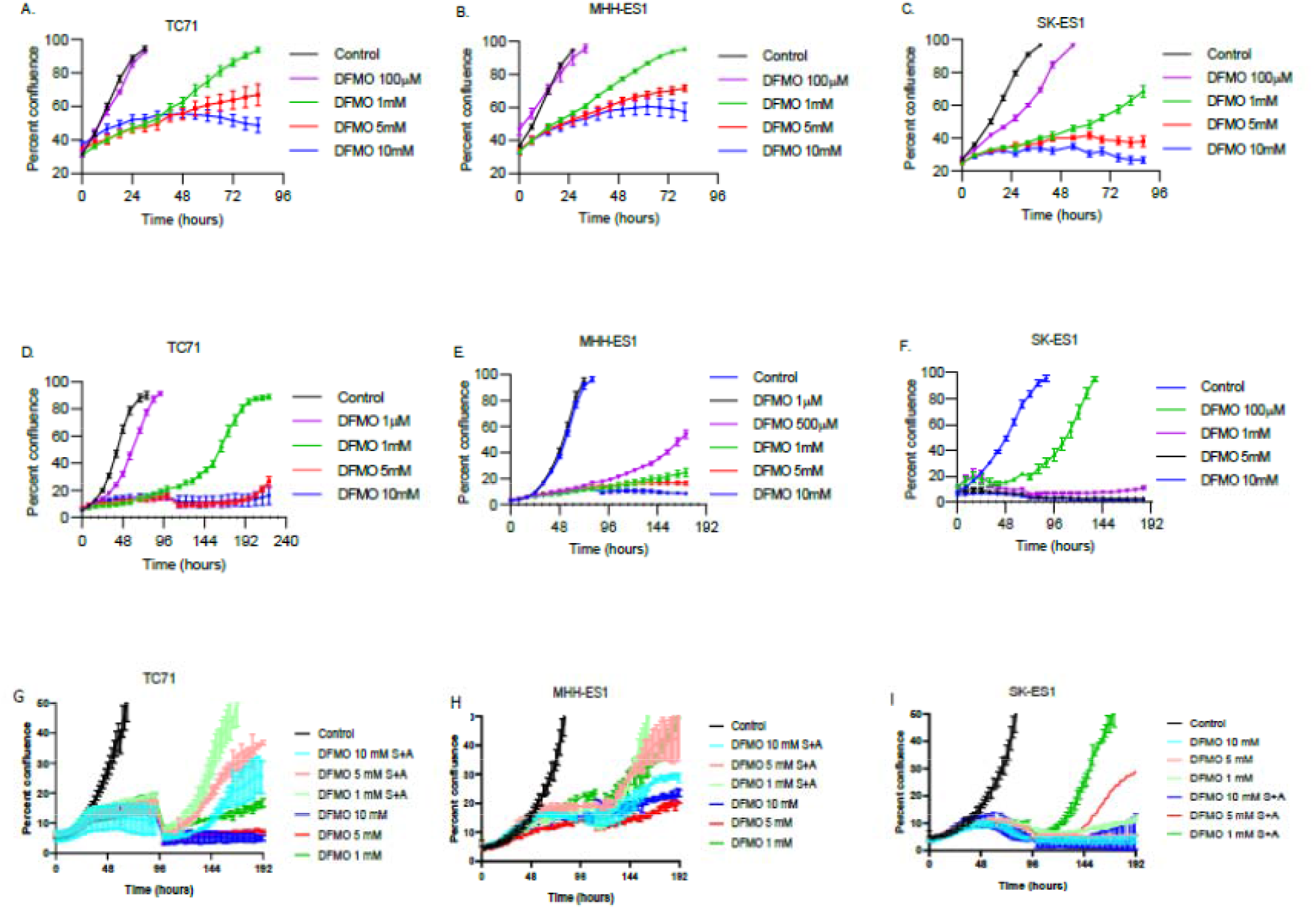
DFMO inhibits Ewing sarcoma proliferation in vitro. (A) TC71 (B) SK-ES1 (C) MHH-ES1 cells were treated with DFMO at various concentrations. Error bars represent 95% confidence interval for triplicate evaluation of each sample. This data is representative of 3 independent experiments demonstrating a dose-dependent inhibition of proliferation with exposure to DFMO. (D) TC71 (E) SK-ES1 (F) MHH-ES1 cells were treated with the indicated concentrations of DFMO. After 4 days, DFMO was removed and replaced with fresh media without DFMO. Resumption of proliferation was observed in all 3-cell lines. (G) TC71 (H) SK-ES1 (I) MHH-ES1 cells were treated with the indicated concentrations of DFMO. After 4 days, fresh DFMO was added with or without the addition of spermidine (5 μM) and aminoguanidine (1 mM). Proliferation was measured over an additional 6 days utilizing the Incucyte® Live-Cell Analysis System, and cells were noted to proliferate in cultures supplemented with spermidine and aminoguanidine.

We next investigated where in the cell cycle polyamine-depleted cells arrest. Cells were cultured with or without DFMO at the IC50 established in the proliferation experiments, and cell cycle distribution was monitored by flow cytometry. In all 3 Ewing sarcoma cell lines evaluated, DFMO induced a cell cycle arrest characterized by accumulation of cells in G1 with a concurrent decrease in the proportion of cells in S phase (Figure 3A-F). Taken all together, our data demonstrate that polyamine depletion caused by treatment with DFMO induces a reversible G1 cell cycle arrest in Ewing sarcoma cells *in vitro*. To confirm that DFMO does not induce apoptosis, Ewing sarcoma cells were assayed for activation of caspase 3, a key mediator of this mode of programmed cell death. Whereas etoposide treatment results in profound activation of caspase 3, this was not detected in any of the 3 cell lines tested, even at the highest doses of DFMO (Figure 3G-I).

**Figure 3:**
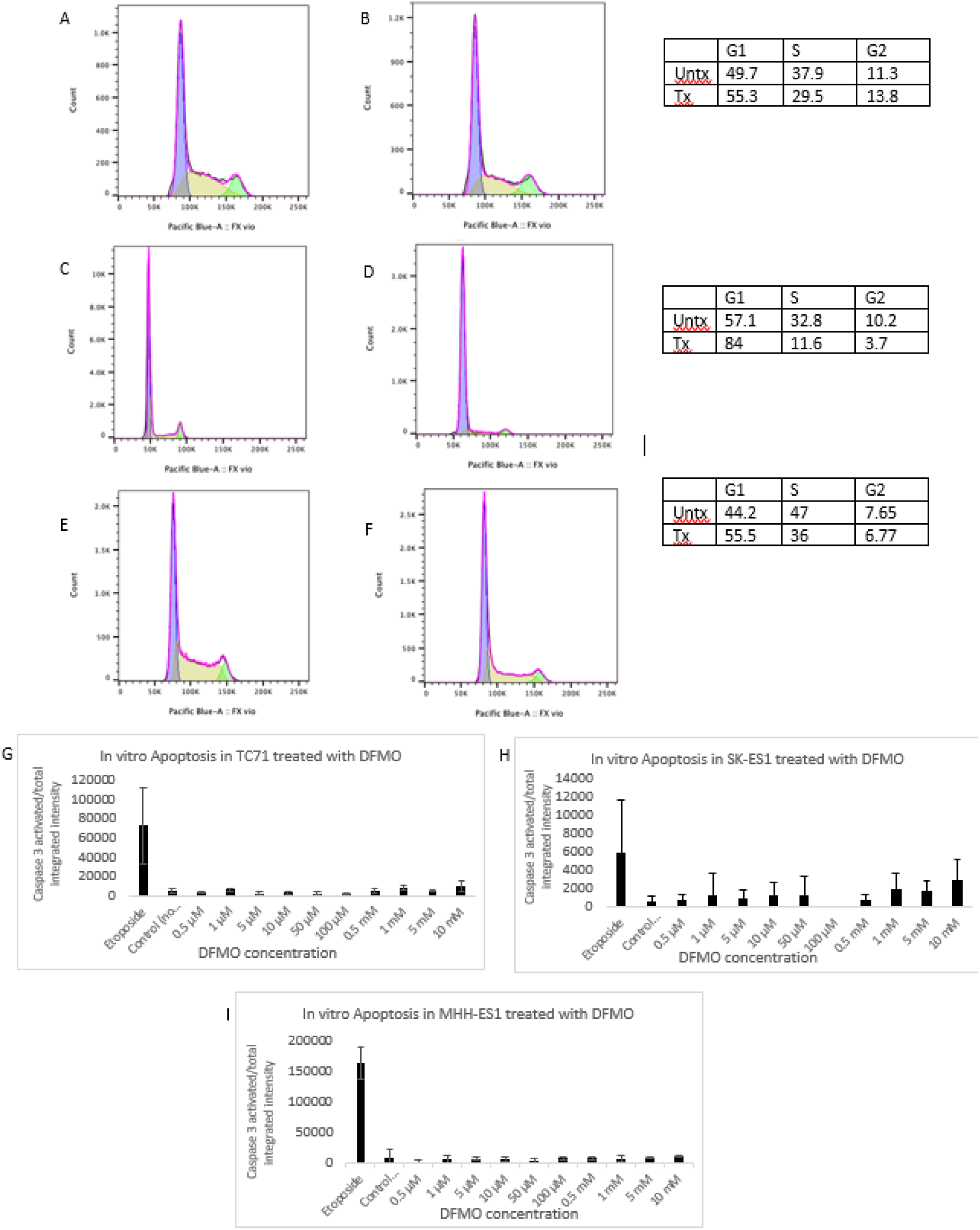
DFMO induces a cell cycle arrest in Ewing sarcoma cells and does not induce apoptosis. Histograms demonstrate the percentage of cells in the G1 (purple), S (yellow), and G2/M (green) phases of the cell cycle. Representative results of experiments performed 2 separate times are shown. The cell cycle arrest is characterized by accumulation of cells in G1 with a concurrent decrease in the proportion of cells in S phase. (A) TC71 cells untreated (B) TC71 cells treated with DFMO at the IC50 (C) SK-ES1 cells untreated (D) SK-ES1 treated with DFMO at the IC50 (E) MHH-ES1 cells untreated (F) MHH-ES1 cells treated with DFMO at the IC50. 300,000 (G) TC-71 (H) SK-ES1 and (I) MHH-ES1 cells were plated in RPMI + 10% FBS. After 24 hours, the media was removed, and fresh media supplemented with Incucyte® Caspase-3/7 Dye for Apoptosis was added. DFMO was added at the indicated concentrations of 0.5 μM to 10 mM. Etoposide (3 μM) was used as a positive control. Plates were inserted into the Incucyte® Live-Cell Analysis System and caspase activity was measured utilizing live-cell time-lapse imaging at 24 hours. Error bars represent 95% confidence interval for triplicate evaluation of each sample. This data is representative of 3 independent experiments. Whereas etoposide treatment results in profound activation of caspase 3 in all 3 cell lines tested, this was not detected in cells treated with DFMO, even at the highest doses tested.

Ewing sarcoma cells with a stem cell phenotype are hypothesized to be the cells that mediate metastasis and chemotherapy resistance (26). The ability to form spherical colonies under non-adherent conditions (sarcospheres) is a well-established surrogate marker of a cancer stem cell phenotype (27-29). To determine if ODC inhibition affects Ewing sarcoma stem cell activity, we investigated the effect of DFMO on the ability of three Ewing sarcoma cell lines to form sarcospheres under non-adherent conditions. MHH-ES1, TC-71, and SK-ES1 cells were plated at clonogenic density on ultra-low adherence plates with varying concentrations of DFMO. Absolute quantification was hindered because the spheres adhere to each other, but after 5 days of growth, we observed a dose-dependent inhibition of sarcosphere growth by DFMO in all 3 cell lines (Figure 4A). Interestingly, DFMO had no impact when added to cultures containing already-established sarcospheres (Figure 4B). Finally, we confirmed that the effect of DFMO on sarcosphere formation is a direct consequence of polyamine depletion by demonstrating that cells cultured in the presence of exogenous spermidine (and aminoguanidine) form sarcospheres despite treatment with DFMO (Figure 4C).

**Figure 4:**
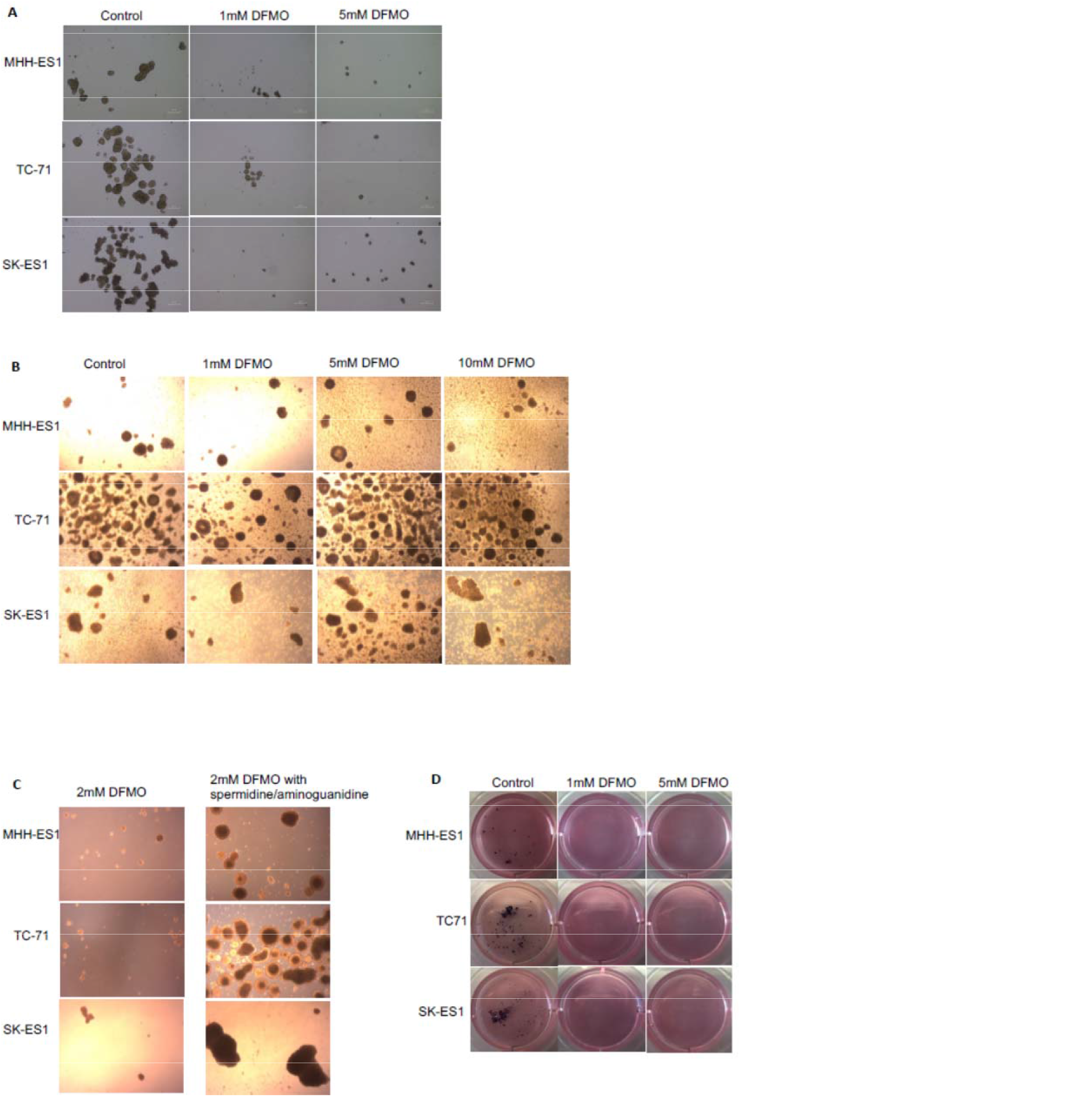
Sarcosphere formation and clonogenic growth in soft agar are inhibited by DFMO. (A) MHH-ES1, TC-71, and SK-ES1 were grown in non-adherent conditions with non-differentiating media and treated with either 1mM or 5mM DFMO. Images were taken at 40x magnification after 5 days of growth. (B) MHH-ES1, TC-71, and SK-ES1 were grown in non-adherent conditions with non-differentiating media. After allowing for spheroids to form, DFMO was added at either 1mM, 5mM or 10mM. Images were taken at 40x magnification after 5 days of growth. (C) MHH-ES1, TC-71, and SK-ES1 were grown in non-adherent conditions with non-differentiating media and treated with either 2mM DFMO or 2mM DFMO supplemented with spermidine and aminoguanidine. Images were taken at 40x magnification after 5 days of growth. (D) Single cells from the MHH-ES1, TC-71, and SK-ES1 cell lines were suspended in soft agar and treated with either 1 mM or 5 mM DFMO. Their growth was monitored for three weeks before the colonies were fixed and stained with p-Nitro Blue Tetrazolium Chloride. No colonies were grossly observed at either treatment concentration, nor were any apparent under microscopic examination.

Colony formation in soft agar is a second established assay to evaluate for the cancer stem cell phenotype and clonogenic potential (30). MHH-ES1, TC-71, and SK-ES1 cells were plated at low density in a soft agar matrix with or without increasing concentrations of DFMO and cultured for three weeks. DFMO at 1 mM completely eliminated soft agar colony growth in all tested cell lines (Figure 4D). Taken together, these data suggest that DFMO inhibits Ewing sarcoma stem cell activity as a direct result of polyamine depletion, supporting the potential use of this agent to prevent Ewing sarcoma growth and metastasis *in vivo*.

### DFMO slows Ewing sarcoma tumor growth in vivo, prolongs survival, and inhibits metastasis

Informed by our *in vitro* results indicating DFMO inhibited the proliferation of Ewing sarcoma cells in culture and also eradicated both sarcosphere growth and soft agar colony formation in multiple Ewing sarcoma cell lines, we investigated the effect of DFMO on Ewing sarcoma tumor growth and metastasis *in vivo*. To determine whether DFMO had the capacity to inhibit tumorigenicity in vivo, 100 TC-71 cells were injected subcutaneously into NOD/SCID/IL-2Rγ-null (NSG) mice. The mice were randomized to receive either 1% DFMO (w/v) treated water or untreated water beginning the day of implantation. The mice were followed closely and there was no significant difference in tumor initiation, as both treated and untreated mice developed tumors nearly uniformly (9/10 control mice and 10/10 DFMO-treated mice developed tumors) and over a similar timeframe. The experiment was then repeated with 2% DFMO (w/v) instead of 1%. In this experiment, 20/20 control mice developed tumors that grew to a diameter of 1.5 cm within 8 weeks, whereas only one of the 20 mice provided with DFMO-supplemented water developed any tumors during that timeframe. Thus, 2% DFMO prevents tumor initiation when a limiting number of Ewing sarcoma cells are implanted in immune deficient mice.

To investigate the effect of DFMO on the growth of established tumors, fragments of a low passage, ES patient-derived xenograft (JHHESX3) were implanted either orthotopically or subcutaneously in NSG mice. After recovery from surgery and confirmation of tumor growth, cohorts of mice were provided with water containing 1% DFMO or with water lacking DFMO. Treatment with 1% DFMO did not affect tumor growth velocity in either tumor location when compared to the control group (Supplemental Figure 1A and 1B). A potential explanation for the observed lack of activity in vivo is that 1% DFMO could be insufficient to exert the inhibitory effects demonstrated *in vitro*, possibly due to rapid excretion in the urine (31) or because of its poor transport profile (32). To address this possibility, the experiment was repeated with a higher DFMO concentration. NSG mice were implanted with fragments of the JHHESX3 PDX in either orthotopic or subcutaneous locations and the cohorts were supplied with untreated water or water treated with 2% DFMO (w/v). One cohort of NSG mice was treated with DFMO immediately upon confirmation of tumor growth (early treatment group), while another cohort was treated post-amputation (adjuvant treatment group). To confirm adequate dosing to inhibit ODC activity, urine was collected from control mice and from the early treatment mice on Day 7 of DFMO treatment, and putrescine and spermidine were quantified. Mice treated with 2% DFMO had a significant reduction in both urinary putrescine (p=0.0001) and spermidine (p=0.024) (Supplemental Figure 2A). The mice were observed until maximal tumor diameter was 1 cm and then pretibial orthotopic tumors were amputated to allow for the development of metastatic disease. The mice with subcutaneous tumors were sacrificed and their tumors collected when maximal tumor diameter reached 1.5 cm. Tumor samples from each cohort were analyzed for ODC activity. As seen previously, subcutaneous tumors expressed higher ODC activity than orthotopic tumors (429.1±67.9 vs 183.6±25.1 pmol CO2/hr/mg protein p=0.014). Orthotopic tumors from DFMO-treated mice demonstrated lower ODC activity than tumors from control mice (204.2±18.1 vs 113.0±22.0 pmol CO2/hr/mg protein, p=0.018), confirming suppression of ODC activity by this dose of DFMO (Supplemental Figure 2B). To confirm that inhibiting ODC results in depletion of intracellular polyamines, we quantified polyamine content in both subcutaneous and orthotopic tumors from mice treated with or without 2% DFMO. Concordant with the urine analysis, putrescine and spermidine levels were significantly reduced in both pretibial and subcutaneous tumors in mice treated with 2% DFMO (Supplemental Figure 2C and 2D).

Spermine levels were not diminished in DMFO-treated tumors, congruent with previous results in DFMO-treated rat hepatoma cells (33).

Having confirmed that 2% DFMO inhibits ODC activity in Ewing sarcoma PDXs, and that this correlates with decreased levels of polyamines, we next evaluated the impact of DFMO on the growth and metastasis of tumors implanted orthotopically. We found that 2% DFMO prolongs the median time to amputation (based on tumor reaching maximum diameter of 10 mm or evidence of animal distress) from 75 days to 92 days, a statistically significant improvement in time to amputation (p=0.0124). Unfortunately, mice treated with DFMO from the time of confirmation of tumor growth suffered excessive perioperative mortality, making it impossible to determine an impact of this early treatment on metastasis. The adjuvant treatment cohort of mice in this experiment was given untreated water during pre-amputation tumor growth and then DFMO-supplemented water after hindlimb amputation. This cohort most accurately mimics the clinical course of a typical Ewing sarcoma patient, who would be expected to have a significant primary tumor in place at the time of diagnosis. In both our orthotopic implantation animal model and in newly diagnosed patients, micrometastatic disease has disseminated by the time of the initiation of treatment. Treatment with 1% DFMO did not prolong survival of mice in either cohort (Supplemental Figure 1C). The mice treated with 1% DFMO died of metastatic disease at nearly identical rates as untreated animals. Mice in the adjuvant treatment cohort were treated with 2% DFMO upon recovery from amputation and were observed for 45 days prior to euthanasia and necropsy to determine the extent of metastatic disease. Although most of the control mice died of metastatic disease during this time, no mice in the cohort treated with 2% DFMO died of metastases, a significant improvement in survival (p=0.0495; Figure 5A). In control mice, metastatic tumors were found within the abdomen including renal and ovarian tumors (Figure 5B) as well as in the lung. These masses were confirmed as Ewing sarcoma by demonstrating expression of human NKX2.2 by immunohistochemistry (Figure 5D and E). Further, there were additional lesions noted, exclusively in the DFMO-treated cohort, that were small (Figure 5C) and necrotic (Figure 5F and G), suggestive of treated metastatic disease with limited NKX2.2 staining. Seven of nine mice in the 2% DFMO-treated cohort were devoid of metastasis, compared with four of nine control mice that lacked metastatic disease. Because we noted differences not only in the quantity but also in the size of metastatic lesions in control mice compared with DFMO-treated mice, we quantified metastasis using a Metastatic Index. Metastatic lesions were assigned a numeric value according to the size of each tumor (small tumors assigned a value of 1, medium tumors a value of 2, and large tumors a value of 3) by a blinded observer. For each mouse, the sum of their assigned tumor values was calculated and termed their Metastatic Index. For example, a mouse with 2 small lesions would be assigned a Metastatic Index of 2, whereas a mouse with 1 small lesion and 2 large lesions would be assigned a Metastatic Index of 7. This approach was designed to mimic the standardized sum of cross products utilized to quantify disease response in pediatric oncology clinical trials. The mean Metastatic Index in the control mice was 1.89, compared with 0.44 in the DFMO-treated mice. This difference was statistically significant (p=0.045; Figure 5H) and demonstrates that, given post-amputation, adjuvant DFMO can inhibit the metastatic outgrowth of disseminated Ewing sarcoma cells.

**Figure 5:**
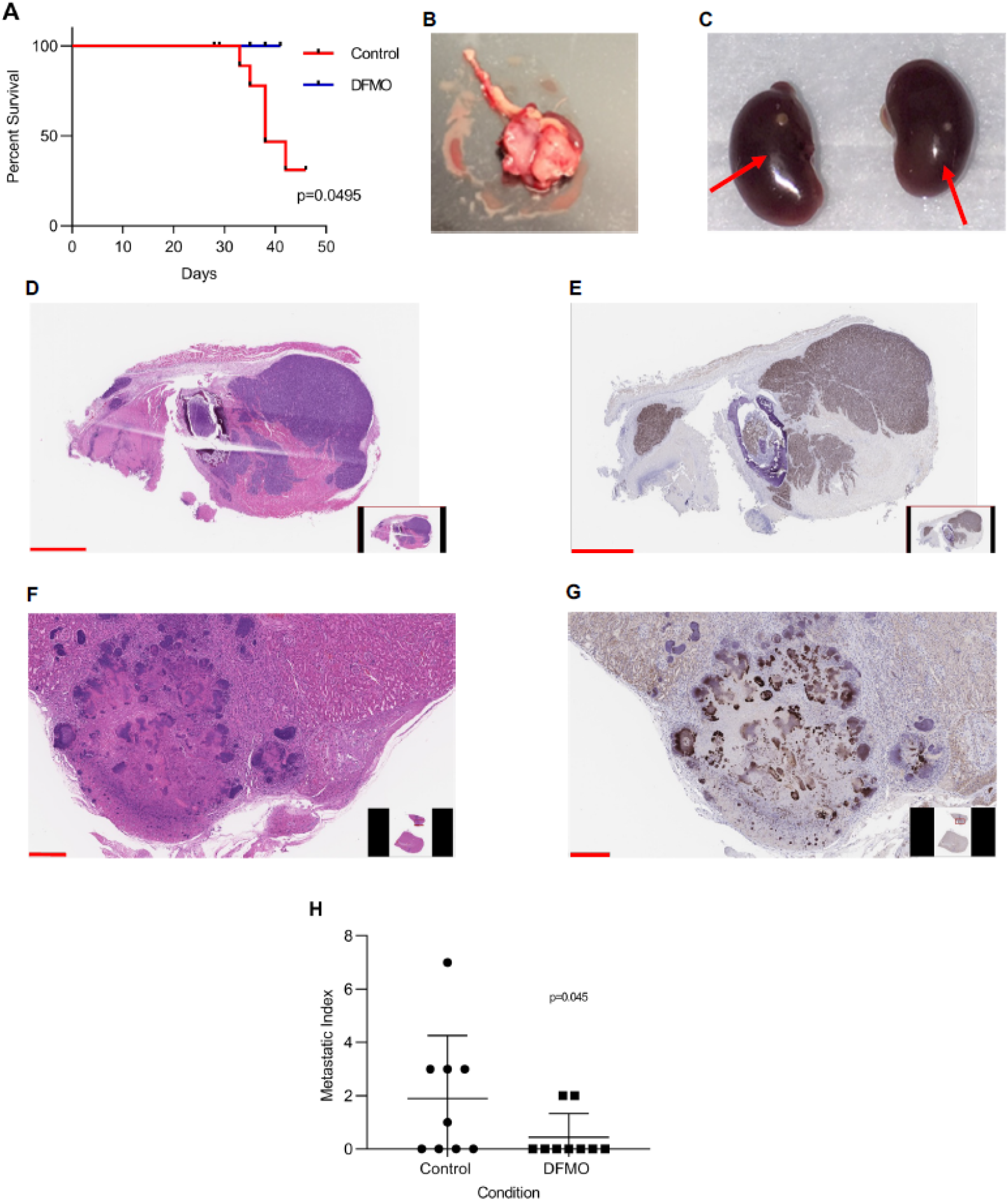
2% DFMO inhibits Ewing sarcoma metastasis and prolongs survival. Mice had fragments of the JHHESX3 PDX implanted orthotopically into the pretibial space and were given either regular drinking water or water supplemented with 2% DFMO. Hindlimbs were amputated when tumor circumference was 30 mm. Panel A shows time to amputation for control mice (n=8) compared with mice treated with 2% DFMO (n=9). All mice were euthanized in the 5th week following amputation and analyzed for metastasis by necropsy. Panel B shows a large renal/ovarian metastasis, and panel C (red arrows) demonstrates very small renal metastases. A lymph node metastasis from a control mouse stained with H&E (D) demonstrates strong NKX2.2 staining by immunohistochemistry (E) (scale bars 2mm). A tiny renal metastasis stained with H&E (F) and evaluated for NKX2.2 by immunohistochemistry(G) appears necrotic (scale bars 200um). Total metastatic burden was evaluated by quantifying Metastatic Index and demonstrates a statistically significant decrease in Metastatic Index in the DFMO-treated mice (H).

Having demonstrated that adjuvant 2% DFMO decreases the metastatic burden in mice with Ewing sarcoma, we next investigated the combination of neoadjuvant ifosfamide followed by adjuvant DFMO. Fragments of the JHHESX3 Ewing sarcoma PDX were implanted in the hind limbs of NSG mice. When tumor growth was established, mice were randomized to treatment with ifosfamide (60 mg/kg/day x 3 days repeated every 3 weeks; 20 mice) or DMSO control (11 mice). Hindlimbs were amputated when tumor diameter was 1.2 cm. Mice treated with neoadjuvant ifosfamide were then randomized to receive either drinking water supplemented with 2% DFMO (w/v) or regular water (10 mice per cohort). Although neoadjuvant ifosfamide did not prolong recurrence-free survival after hind limb amputation, the addition of adjuvant DFMO did prolong survival compared to control mice (Figure 6A). Of note, only 2 mice in the DFMO-treated cohort died of recurrent disease, compared with all of the control mice. We evaluated the burden of metastatic disease in each mouse by necropsy. The mice treated with neoadjuvant ifosfamide followed by adjuvant 2% DFMO had a statistically significant decrease in metastatic index compared with both control mice and those treated solely with neoadjuvant ifosfamide (Figure 6B). Thus, adjuvant DFMO prolongs survival and limits metastatic recurrence both as a single agent and when given to mice previously treated with chemotherapy.

**Figure 6:**
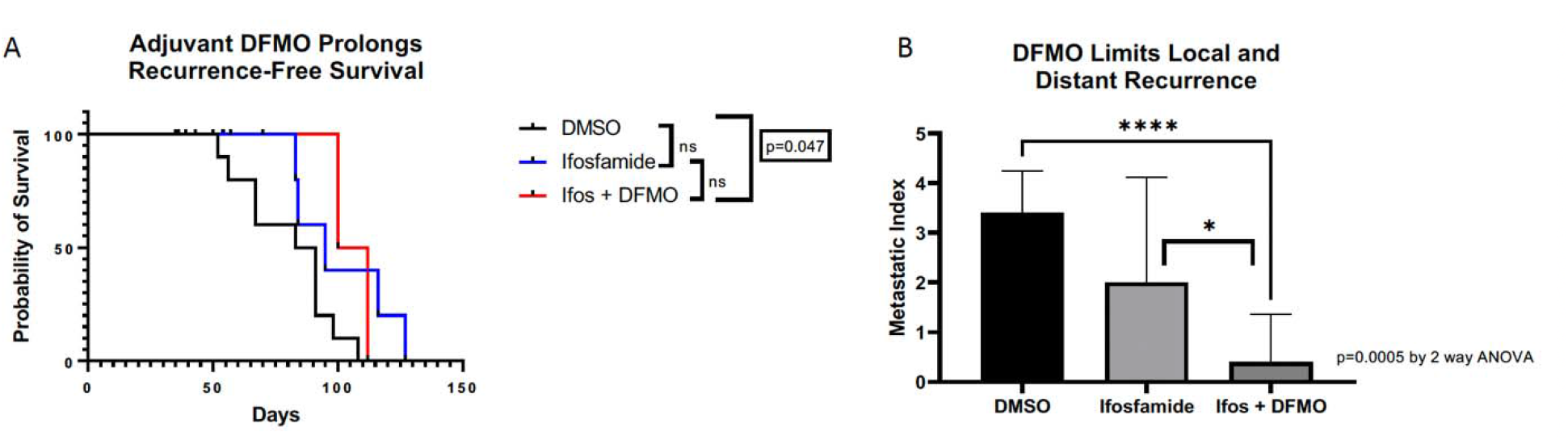
The combination of neoadjuvant ifosfamide and adjuvant DFMO prolongs recurrence-free survival and limits local and distance recurrence. Mice had fragments of the JHHESX3 PDX implanted orthotopically into the pretibial space and were given either DMSO or Ifosfamide upon tumor engraftment. Hindlimbs were amputated when tumor diameter was 12 mm. Mice treated with Ifosfamide were then randomized to receive either 2% DFMO or regular drinking water. (A) Adjuvant DFMO after neoadjuvant ifosfamide prolongs recurrence-free survival from tumor amputation compared with control (p=0.047). (B) Total metastatic burden was evaluated by quantifying Metastatic Index and demonstrates a statistically significant decrease in Metastatic Index in the Ifosfamide + DFMO-treated mice as compared to both the DMSO treated mice and Ifosfamide alone (p=0.0005 by 2-way ANOVA)).

### DFMO primes cells for ferroptosis in vitro

Because the small metastatic nodules noted in DFMO-treated mice are necrotic upon histological examination (Figure 5F and G), but DFMO does not induce apoptosis *in vitro* in cells growing in 2D culture, we investigated other mechanisms by which DFMO might kill disseminated tumor cells. There is increasing recognition that ferroptosis is an important restraint on metastasis (34-36), and it was recently reported that p53-mediated induction of ferroptosis requires expression of spermidine/spermine N1-acetyltransferase 1 (SAT1), the rate limiting catabolic enzyme in polyamine excretion from cells. This group also reported that induction of SAT1 is correlated with increased expression of arachidonate 15-lipoxygenase (ALOX15), a key mediator of the lipid peroxidation that characterizes ferroptosis (37). Based on these findings, we investigated the impact of DFMO on ALOX15 expression in Ewing sarcoma cells *in vitro*. We found that treatment with DFMO induces a statistically significant increased expression of ALOX15 in 2 of the 3 Ewing sarcoma lines we tested (Figure 7A). This increase is abolished if cells are concurrently provided exogenous spermidine (and aminoguanidine; Figure 7B), consistent with the increase in ALOX15 expression being a direct result of polyamine depletion. Because ferroptosis can be inhibited by cell-cell and cell-matrix contact (40), we investigated whether the inhibition of sphere formation by DFMO (Figure 4) reflects induction of ferroptosis in the single cell suspension that is plated onto the nonadherent tissue culture plastic. MHH-ES1, TC-71, and SK-ES1 cells were plated at clonogenic density on ultra-low adherence plates with varying concentrations of ferrostatin-1 in the presence of DFMO. After 7 days of growth, we observed well-established sarcospheres in a dose dependent manner in all 3 cell lines (Figure 7C). Thus, *in vitro*, polyamine depletion results in increased expression of ALOX15, a key executioner of ferroptosis, and this induction of ferroptosis is the mechanism underlying inhibition of sphere formation by DFMO.

**Figure 7:**
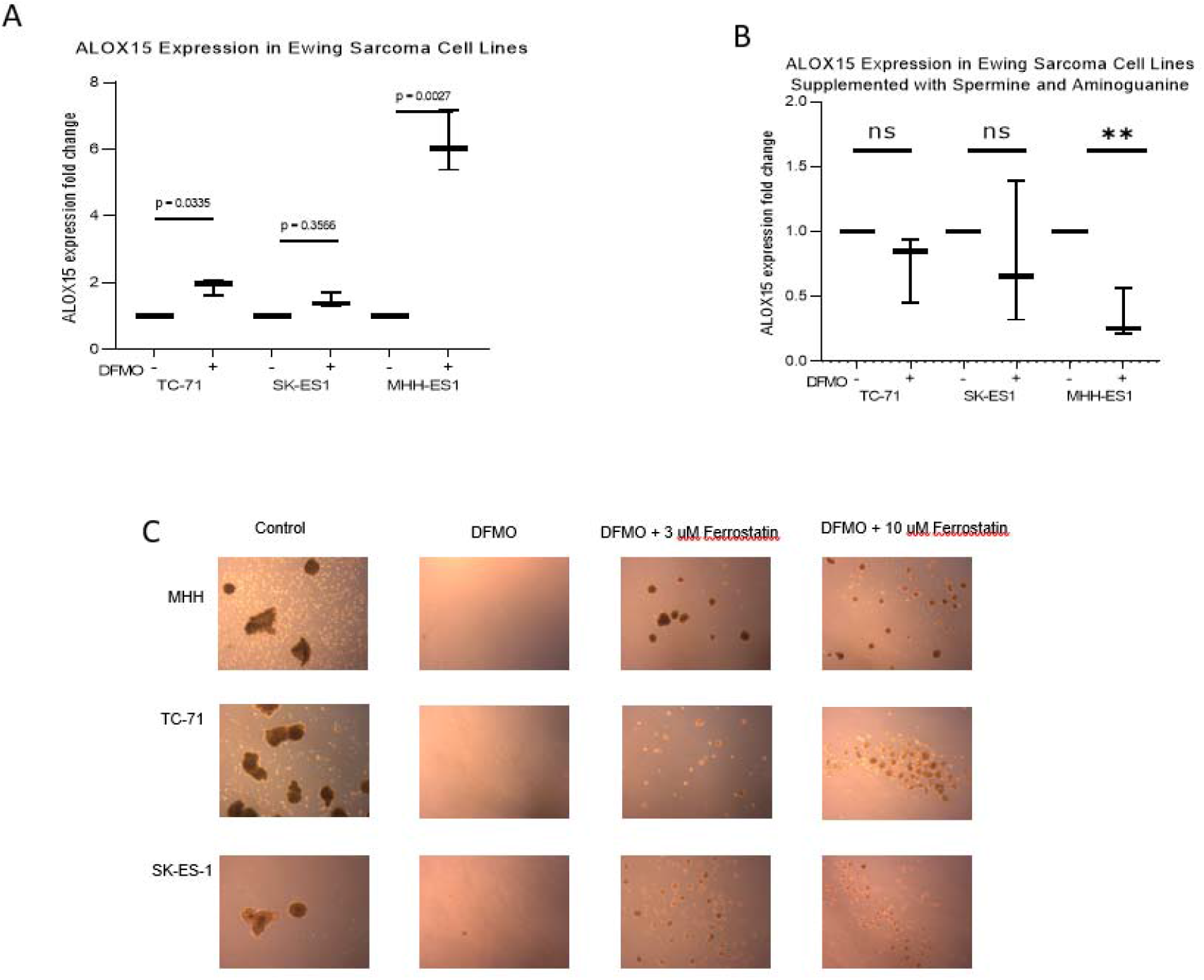
DFMO induces ALOX15 expression. (A) TC-71, MHH-ES1 and SK-ES1 cells were plated per well and DFMO was added at the IC50 previously determined for each cell line, with and without spermidine and aminoguanidine. ALOX15 mRNA expression was quantified by qRT-PCR. We found that treatment with DFMO induces a statistically significant increased expression of ALOX15 in 2 of the 3 Ewing sarcoma lines tested. This increase is abolished if cells are concurrently provided exogenous spermidine (and aminoguanidine) (B). Representative results are shown. Experiment was repeated 3 times with similar results. Error bars represent standard error of the mean. (C) MHH-ES1, TC-71, and SK-ES1 were grown in non-adherent conditions with non-differentiating media and treated with either DFMO or DFMO supplemented with 3uM Ferrostatin or 10uM Ferrostatin. Images were taken at 40x magnification after 7 days of growth.

### Disseminated Ewing sarcoma cells display an increased vulnerability to ferroptosis which is triggered by DFMO

Next, we investigated the ability of DFMO to induce ferroptosis *in vivo*. First, we identified gene expression differences between disseminated Ewing sarcoma cells and the primary tumor. To accomplish this, we implanted tumors composed of TdTomato-labeled TC71 cells in the tibias of NSG mice. When tumors reached 1.5 cm in diameter, Ewing sarcoma cells were recovered from lungs, bone marrow, and the primary tumor by flow cytometry. RNA was isolated from these cells and analyzed by RNA sequencing. Analysis of the expression of the top 500 most differentially expressed genes demonstrated that, at the transcriptomic level, Ewing sarcoma cells collected from the bone marrow were more similar to those from the primary tumor when compared to the Ewing sarcoma cells collected from lungs (Figure 8A) which showed a distinct transcriptomic profile. KEGG analysis of differentially expressed genes showed that cells isolated from the lungs had downregulation of genes involved in both glutathione metabolism and the pentose phosphate pathway. This finding was more thoroughly evaluated by examining expression levels of a panel of genes involved in glutathione metabolism and the pentose phosphate pathway (Figure 8B). Glutathione is the major system employed by cells to counter the accumulation of reactive oxygen species which contribute to lipid peroxidation. For glutathione to be recycled and maintain a reduced environment requires NADPH, and the major source of this metabolite is the pentose phosphate pathway. We also interrogated the expression of a panel of genes associated with ferroptosis and found decreased expression of acyl-CoA Synthetase Long Chain Family Member 3 (ACSL3) and increased expression of Prostaglandin-Endoperoxide Synthase 2 (PTGS2) (Supplemental Figure 3A). ACSL3 activates long-chain fatty acids for beta-oxidation, so decreased ACSL3 would favor the accumulation of long-chain fatty acids, which are peroxidated to trigger ferroptosis. PTGS2 is a lipid cyclooxygenase and peroxidase, and its cyclooxygenase activity oxygenates arachidonate to the hydroperoxy endoperoxide prostaglandin G2. The net effect of these changes is to downregulate systems that would oppose ferroptosis (glutathione metabolism, the pentose phosphate pathway, and ACSL3) and to upregulate a lipid peroxidase, all of which are changes that would increase the vulnerability of these cells to ferroptosis.

**Figure 8:**
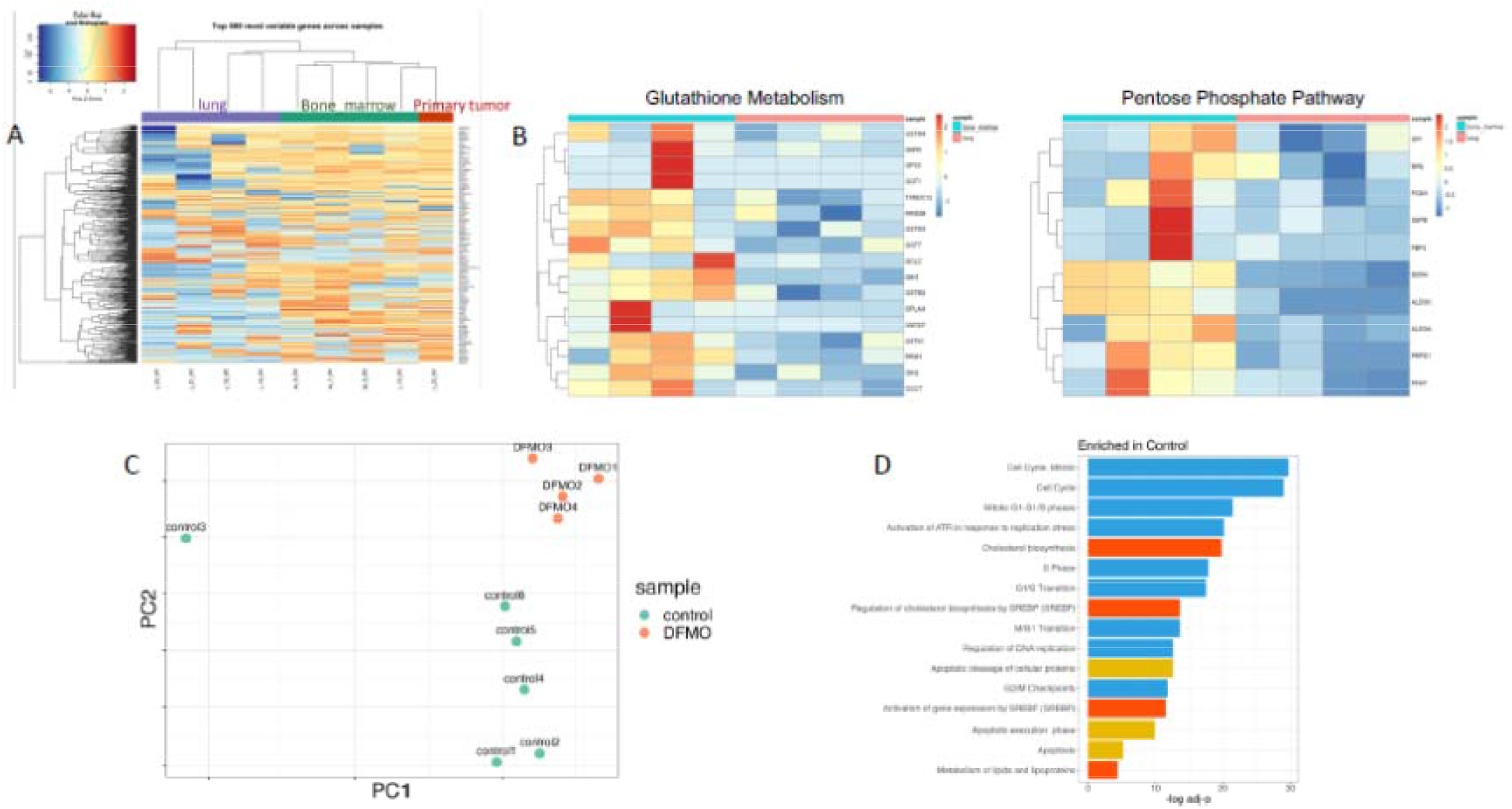
Transcriptomic analysis demonstrates that disseminated tumor cells are vulnerable to, and that DFMO induces, ferroptosis. To identify gene expression differences between disseminated Ewing sarcoma cells and the primary tumor, tumors composed of TdTomato-labeled TC71 cells were implanted in the tibias of NSG mice. When tumors reached 1.5 cm in diameter, Ewing sarcoma cells were recovered from lungs, bone marrow, and the primary tumor by flow cytometry. RNA was isolated from these cells and analyzed by RNA sequencing. (A) Unsupervised clustering of samples based on gene expression changes demonstrated that, at the transcriptomic level, Ewing sarcoma cells collected from the bone marrow were more similar to those from the primary tumor when compared to the Ewing sarcoma cells collected from lungs (B) Enrichment analysis of differentially expressed genes showed that cells isolated from the lungs had downregulation of genes involved in both glutathione metabolism and the pentose phosphate pathway. To identify transcriptomic changes induced by DFMO, we performed RNA sequencing of tumors grown in the tibias of NSG mice treated with or without DFMO (2% w/v) for 14 days. (C) Principal Component Analysis demonstrated clustering of the tumors from control mice separate from tumors exposed to DFMO (D) Enrichment analysis of differentially expressed genes demonstrated downregulation of genes associated with cell cycle progression, induction of apoptosis, and cholesterol biosynthesis in tumors collected from DFMO-treated mice compared with control.

We next investigated the impact of DFMO. Because disseminated Ewing sarcoma cells are killed by DFMO (Figure 5F), we could not isolate DFMO-treated disseminated cells for analysis. We therefore performed transcriptomic analysis of Ewing sarcoma PDXs, implanted orthotopically in NSG mice provided with either plain drinking water or water supplemented with 2% DFMO. Principal Component Analysis demonstrated clustering of the tumors from control mice separate from tumors exposed to DFMO (Figure 8C). Analysis of differentially expressed genes demonstrated downregulation of genes associated with cell cycle progression, induction of apoptosis, and cholesterol biosynthesis (Figure 8D). Querying a list of genes associated with ferroptosis (Supplemental Figure 3B) revealed downregulation of Heat Shock Protein Family B Member 1 (HSPB1) and 3-hydroxy-3-methylglutaryl-CoA reductase (HMGCR) and upregulation of lysophosphatidylcholine acyltransferase 3 (LPCAT3). HSPB1 is a negative regulator of ferroptosis (38). HMGCR is the rate-limiting step in cholesterol biosynthesis and downregulation of this enzyme results in decreased levels of CoQ10, an inhibitor of lipid peroxidation (39). LPCAT3 catalyzes the esterification reaction that inserts arachidonic acid into phospholipids, a necessary early step in priming cells for ferroptosis (39). These gene expression changes, especially in the context of the gene expression changes induced through the process of dissemination, support our model that DFMO inhibits metastatic outgrowth of disseminated Ewing sarcoma cells by inducing ferroptosis.

## Discussion

Previous work from our laboratory demonstrated that arginase is differentially expressed in fragments of Ewing sarcoma xenograft implanted in the tibia (orthotopically) of immune deficient mice compared to fragments implanted subcutaneously (6). Because tumor fragments implanted orthotopically undergo spontaneous distant metastasis while tumor fragments implanted subcutaneously do not, and because arginase catalyzes the production of ornithine, which can be converted into putresceine by ODC, we investigated whether polyamine synthesis plays a role in Ewing sarcoma metastasis. We found that DFMO, an irreversible inhibitor of ODC, is cytostatic to Ewing sarcoma cells growing in culture and prevents both clonogenic growth in soft agar and growth as sarcospheres under nonadherent conditions. We further found that 2% DFMO-supplemented drinking water slows the growth of an orthotopically implanted Ewing sarcoma PDX and substantially inhibits the development of metastases.

Our data support the model that disseminated Ewing sarcoma tumor cells are sensitized to undergo ferroptosis, and that the polyamine depletion induced by treatment with DFMO triggers this form of programmed cell death, preventing the disseminated cells from becoming macrometastases. Recently 3D cell culture methods have been integrated into cancer research as they more closely replicate many aspects of *in vivo* tumors. Our findings using 2D culture of Ewing sarcoma cells, that DFMO induces a reversible cell cycle arrest and does not induce apoptosis, did not jibe with our findings *in vivo* that DFMO prevented the metastatic outgrowth of disseminated tumor cells, and that whatever small lesions formed were necrotic at the time of necropsy. Thus, we turned to a 3D cell culture technique, the ability to grow as spheres under nonadherent conditions, and were able to demonstrate that DFMO induces ferroptosis in this system. In our *in vivo* models, we found that disseminated Ewing sarcoma cells have transcriptional changes that include downregulation of glutathione metabolism and the pentose phosphate pathway, and that polyamine depletion by DFMO results in a second series of gene expression changes that should combine to trigger a significant increase in peroxidated lipids, triggering ferroptosis. Supportive of this model is our finding that DFMO prevents the formation of sarcospheres when cells are plated under nonadherent conditions in vitro, but that DFMO does not impact previously formed sarcospheres. This finding is in line with published data showing that cell-cell contact, such as is seen in sarcospheres, inhibits ferroptosis (40). Recently, Zhang et al. (41) reported that IL1RAP is a key driver of Ewing sarcoma metastasis through its ability to help cells overcome the oxidative stress of anoikis by maintaining cysteine and glutathione pools. Our findings can be reconciled with theirs because a) we see an effect on cells that have already disseminated, so anoikis is not a barrier to growth of microscopic metastases into macroscopic nodules, b) the specific genes we see altered in disseminated cells related to glutathione are transferases (GSTM1 and GSTP1, for example) that would prevent utilization of glutathione regardless of concentration, c) decrease in flux through the pentose phosphate pathway would be expected to diminish intracellular NADPH, which is necessary for glutathione recycling but would not alter cysteine levels, and d) if production of peroxidated lipids outpaces their reduction by glutathione, they might still accumulate to a level high enough to drive ferroptosis despite the function of IL1RAP. Taken together, our results provide a mechanistic explanation for the observation that adjuvant treatment with DFMO can prevent Ewing sarcoma metastasis in a mouse model and reveal a previously unrecognized function for polyamines – prevention of ferroptosis.

Our model does not preclude a direct effect on Ewing sarcoma stem cells as well. Several of our experiments are consistent with a stem-cell specific effect of DFMO. DFMO prevented both sphere formation and clonogenic growth in soft agar, both of which are in vitro surrogates for a cancer stem cell phenotype, and DFMO (2% w/v) in the drinking water prevented tumor growth when a limiting number of cells was implanted subcutaneously, the gold standard *in vivo* assay of cancer stem cell phenotype. This is in stark contrast to the lack of significant impact of DFMO on growth of established tumors. The mechanism by which DFMO inhibits Ewing sarcoma stem cell function has not yet defined. It could be through induction of ferroptosis, as in disseminated tumor cells, but could also be via a distinct mechanism unique to this rare cell population. Further work will be necessary to define this mechanism.

Although never investigated in Ewing sarcoma, DFMO has been studied as both a chemopreventive agent and as a chemotherapeutic agent in a variety of tumor types (19-22). Importantly from a translational perspective, DFMO shows promise as a maintenance therapy in clinical trials in patients with high risk neuroblastoma (23) and anaplastic gliomas (42). Our results suggest the potential for a similar role for patients with Ewing sarcoma. The data presented here show that adjuvant treatment with DFMO (after tumor cells have already disseminated) of mice bearing orthotopic tumors prevents the outgrowth of metastases and may be cytotoxic in this setting. We also present evidence that adjuvant DFMO augments the impact of neoadjuvant ifosfamide in preventing disease recurrence and prolonging survival. Given the substantial evidence of safety and tolerability of DFMO in both children and adults, this finding supports aggressive study in clinical trials of patients at high risk for metastatic relapse. Our finding that DFMO decreases polyamine levels in the urine of tumor-bearing mice also suggests the possibility that these trials could include a pharmacodynamic component, with ongoing measurement of urinary polyamines ensuring that DFMO dosing is adequate to inhibit ODC. In preclinical studies of neuroblastoma, DFMO had minimal impact as a single agent, presumably because the inhibition of polyamine synthesis can be overcome by import of extracellular polyamines, either by solute carrier family 3 member 2 (SLC3A2) as proposed by Gamble et al. or by other mechanisms (43). An important point for the translational relevance of our findings is that DFMO <u>as a single agent</u> dramatically reduced Ewing sarcoma metastasis, which increases the likelihood of successful translation of our findings from the laboratory to the clinic.

In summary, we found that adjuvant administration of DFMO to mice bearing Ewing sarcoma xenografts prevents the metastatic outgrowth of disseminated tumor cells. Our data support a model in which disseminated Ewing sarcoma cells are uniquely vulnerable to ferroptosis, and this form of programmed cell death is triggered by treatment with DFMO. This is the first direct evidence that intracellular polyamines protect cells from ferroptosis, and future work will investigate whether this is unique to Ewing sarcoma or a more widespread phenomenon, which would suggest that DFMO could be widely used to prevent disseminated tumor cells from becoming gross metastatic disease in multiple cancer types.

## MATERIALS AND METHODS

### Cell lines and xenografts

Established Ewing sarcoma cell lines TC-71, MHH-ES, and SK-ES-1, originally purchased from ATCC, were cultured in 125 cm^2^ tissue culture flasks supplemented with RPMI-1640 (Gibco) + 10% FBS and maintained under standard conditions. Cell line authentication was performed using STR analysis by the Genomics Core at the Albert Einstein College of Medicine. Cells were counted by Trypan blue exclusion (Gibco) and by automated cell counter (TC20, BioRad). We utilized two Ewing sarcoma patient-derived xenografts (PDXs) for *in vivo* experiments; EWS4 (generous gift from Chand Khanna, National Institutes of Health) and JHHESX3, a xenograft established within our lab from a relapsed tumor after obtaining informed consent under a protocol approved by the Johns Hopkins University Institutional Review Board. Xenografts were grown in NOD/SCID/IL-2Rγ null (NSG) mice originally obtained from Jackson Laboratories and then bred by our group. All mouse procedures were performed according to protocols approved by the Johns Hopkins Animal Care and Use Committee and the Institutional Animal Care and Use Committee at Albert Einstein College of Medicine.

### *In vivo* studies

TC-71 cells were cultured, and 100 cells were suspended in Matrigel 1:1 dilution with phosphate-buffered saline in a total volume of 0.1 ml. This suspension was injected into the subcutaneous space of the flank of NSG mice using a 25-gauge needle. Orthotopic and subcutaneous PDX implantations of 5-6 mm^3^ fragments of donor PDX tumor were performed as previously described, as was hindlimb amputation (6), in mice aged 6-8 weeks. To ensure that results were not biased by the sex of the mouse, each experiment contained equal numbers of males and females. For murine low dose DFMO treatment, a 1% solution of drug dissolved in ddH_2_0 (w/v) and was sterilized through a 0.22 μm filter. High dose DFMO was similarly prepared at 2% (w/v) DFMO. The mice were provided with a full water bottle and allowed access ad lib to treated water exclusively. In some experiments, mice were treated with ifosfamide. This drug was delivered by ip injection at a dose of 60 mg/kg/day for 3 consecutive days every 21 days until tumor was large enough for amputation. Ifosfamide-treated mice were also given ondansetron subcutaneously at a dose of 3 mg/kg prior to receiving each dose of ifosfamide.

Tumor size was measured twice weekly beginning when the tumor was easily palpable, typically at 5 mm in diameter. Subcutaneous tumors were measured using calipers in orthogonal directions and volume estimated with the formula a * b^2^/2, where “a” is the diameter of the long axis of the elliptical tumor and “b” is the diameter of the orthogonal axis in the same plane. The measurement metric of pretibial tumors was limb circumference, estimated using an elliptical formula from hindlimb caliper measurements of 2π√((a^2^ + b^2^)/2), with “a” and “b” as above. In both cases, mice were sacrificed or considered for survival amputation when tumors approached 15 mm in the largest dimension or when animals showed signs of distress, such as a body condition score <2.

To evaluate for metastases, mice were sacrificed in the fifth week following hindlimb amputation. Necropsy was performed and visible metastatic lesions were identified, collected and given a volumetric assessment blindly assigning metastases values of 1, 2, or 3 for small, medium, large tumors. These values were summed for each mouse and assigned as their metastatic index value.

### Immunohistochemistry

Immunohistochemistry was performed by Histoserv (Germantown, MD). Slides were deparaffinized and hydrated through graded alcohols to distilled water, followed by antigen retrieval by boiling in a Tris-EDTA buffer (pH 9). The slides were then blocked with hydrogen peroxide and incubated with the primary antibody (NKX2.2 used at 1:1,000). After rinsing, an undiluted HRP-conjugated anti-rabbit secondary was applied. Following application of the secondary, the IHC staining was developed using DAB, intensified with an enhancing reagent, and counterstained with Hematoxylin. All of the incubations were carried at room temperature, with TBST used as the washing buffer.

### Polyamine content and ODC activity analysis

Polyamine content of cells, tissues (3 subcutaneous and 3 orthotopic JHHESX3 PDXs), and urine were determined by the methods of Kabra, et al. (44). For ODC activity, the preserved tissue was thawed, and enzymatic activity was quantified following the protocol detailed in Seely and Pegg (45). One unit of enzyme activity is defined as the amount of enzyme releasing 1 mmol of ^14^CO_2_ per hour at 37° C. Polyamine concentrations and ODC activity levels are presented as per unit protein, which was determined based on the method of Bradford (Bio-Rad Protein Assay, Hercules, CA).

### *In vitro* DFMO proliferation and washout assays

Two thousand five hundred TC-71, MHH-ES1 and SK-ES-1 cells were plated per well in a 96-well, tissue culture-treated plate (Corning) with 75 uL of RPMI-1640 (Gibco) + 10% FBS and maintained under standard conditions. After 24 hours to allow for adherence, DFMO was added at various concentrations from 0.5 μM to 10 mM. Plates were inserted into the Incucyte^®^ Live-Cell Analysis System and confluence was measured utilizing live-cell time-lapse imaging. To further investigate the cytotoxic vs. cytostatic effect of DFMO, washout assays were performed in the Incucyte^®^ Live-Cell Analysis System. One thousand (to minimize overgrowth) TC-71, MHH-ES1 and SK-ES-1 cells were plated per well in a 96-well, tissue culture-treated plate (Corning) with 75 μL of RPMI-1640 (Gibco) + 10% FBS and maintained under standard conditions. After 24 hours, DFMO was added at the same concentrations of 0.5 μM to 10 mM. After 4 days, DFMO was removed, and fresh media RPMI-1640 (Gibco) + 10% FBS was added. Confluence was measured over an additional 6 days utilizing the Incucyte^®^ Live-Cell Analysis System. To investigate the effect of polyamines in the setting of DFMO, 2,500 TC-71, MHH-ES1 and SK-ES-1 cells were plated as above. After 24 hours, DFMO was added at the same concentrations of 0.5 μM to 10 mM. After 4 days, DFMO was removed, and fresh DFMO was added with or without the addition of spermidine (5 μM) and aminoguanidine (1 mM). Confluence was measured over an additional 6 days utilizing the Incucyte^®^ Live-Cell Analysis System.

### Caspase 3/7 activation

Two thousand five hundred TC-71, MHH-ES1 and SK-ES-1 cells were plated per well in a 96-well, tissue culture-treated plate (Corning) with 75 μL of RPMI-1640 (Gibco) + 10% FBS and maintained under standard conditions. After 24 hours, the media was removed and fresh media supplemented with Incucyte® Caspase-3/7 Dye for Apoptosis was added. DFMO was added at the same concentrations of 0.5 μM to 10 mM. Etopopside (3 μM) was used as a positive control. Plates were inserted into the Incucyte^®^ Live-Cell Analysis System and caspase activity was measured utilizing live-cell time-lapse imaging at 24 hours.

### Sarcosphere formation

TC-71, MHH-ES, and SK-ES-1 cells were grown and dissociated as above and cells were resuspended in MesenCult MSC basal media (Stemcell Technologies) supplemented with MesenCult MSC Stimulatory Supplement (Human) (Stemcell Technologies). One thousand viable cells/4 ml of supplemented media were plated in each well of a Costar Ultra-Low Attachment 6-well plate (Corning). A working solution of DFMO was diluted in MesenCult MSC media and added to each well of the treated conditions for final concentrations of 1 mM and 5 mM. To evaluate the effect of DFMO on already formed spheroids, TC-71, MHH-ES, and SK-ES-1 cells were grown and dissociated as above and plated in Costar wells as above. After established spheroids were formed, DFMO was added at varying concentrations from 1 mM to 10 mM. To evaluate the effect of DFMO in the presence of polyamines, Ewing sarcoma cells were grown and dissociated as above and plated in Costar wells as above. A working solution of DFMO was diluted in MesenCult MSC media and added to each well of the treated conditions for final concentrations of 1 mM and 5 mM. Spermidine 5 uM and aminoguanidine 1 mM were added to all wells. To evaluate the effect of Ferrostatin-1 in the presence of DFMO, Ewing sarcoma cells were grown and dissociated as above and plated in Costar wells as above. A working solution of DFMO was diluted in MesenCult MSC media and added to each well of the treated conditions for final concentrations of 1 mM and 5 mM. Ferrostatin-1 was added at 3 uM and 10 uM respectively.

### Soft agar colony formation

To test clonogenic activity, MHH-ES, SK-ES-1, and TC71 cell lines were seeded at a density of 2,500 cells per well of a standard Corning 6 well plate. Each cell line was replicated in six wells total. Cells were cultured in 0.3% top agar (Invitrogen) prepared from a 1:1 dilution of 0.6% bottom agar in RPMI-1640 medium supplemented with 10% FBS. After 1 hour, top agar was overlaid with 1 ml fresh media. Cells were maintained in a 37°C tissue culture incubator for 3 weeks. To test the influence of DFMO on colony formation, DFMO was suspended in a working solution of RPMI-1640 and incorporated into both agar layers as well as in the overlying media layer with final concentrations of 1 mM or 5 mM. After 3 weeks, colonies were stained with p-Nitro Blue Tetrazolium Chloride (USB), photographed, and quantified visually.

### Cell cycle analysis by flow cytometry

Five hundred thousand TC-71, MHH-ES1 and SK-ES-1 cells were plated per well in a 6-well, tissue culture-treated plate (Corning) with 3 mL of RPMI-1640 (Gibco) + 10% FBS and maintained under standard conditions. After 24 hours to allow for adherence, DFMO was added at the previously determined IC50. After 72 hours of treatment, flow cytometry was performed utilizing Click-iT™ EdU Cell Proliferation Kit for Imaging, Alexa Fluor™ 647 dye and FxCycle™ Violet stain (F10347).

### Reverse transcriptase polymerase chain reaction

300,000 TC-71, MHH-ES1 and SK-ES-1 cells were plated per well in a 6 well, tissue culture-treated plate (Corning) with 3 mL of RPMI-1640 (Gibco) + 10% FBS and maintained under standard conditions. After 24 hours to allow for adherence, DFMO was added at the appropriate IC50 with and without spermidine and aminoguanidine. RNA was processed via Qiagen RNeasy kit, and converted to cDNA utilizing iscript SuperScript reverse transcriptase RT-PCR was performed using Biorad primer ALOX15.

### RNA sequencing

RNA was isolated using the Qiagen RNeasy kit. Libraries were constructed and sequenced by the Yale Center for Genome Analysis (YCGA). The RNA-seq samples for 1) Ewing sarcoma cells isolated from bone marrow, lung, and primary tumor and 2) DFMO treated and control tumors were aligned using salmon (46)(version 1.9.0). The reference genome (hg38 and mm10 combined) was downloaded from Refgenie (47). Genes with less than 100 reads in more than 3/10 samples were discarded as lowly expressed genes, and only human genes were used for downstream analysis. Differential gene expression analysis was performed using R package DESeq2 (48) (version 1.34.0). Genes with adjusted p-value less than 0.05 and positive logfold change were selected for enrichment analysis using R package 1) FGESA (49) (version 1.26.0) and 2) EnrichR (50) (version 3.1) respectively. The RNA-seq data described in our study are currently being submitted to public repositories. As part of our commitment to data transparency and reproducible science, these data will be made publicly available upon publication.

### Statistical analysis

Statistical comparisons between pooled polyamines and tumor sites as well as ODC activity were made using an unpaired two-tailed Student’s t-test. A p value <0.05 was considered significant. Mice were randomized into treatment groups. All statistical analyses were performed using Prism 6 software (GraphPad Software, Inc.).

## Study Approval

All animal studies described above were reviewed and approved by an appropriate institutional review board and approved by the Institutional Animal Care and Use Committees of Johns Hopkins University School of Medicine (IACUC MO16M139) and Albert Einstein College of Medicine (IACUC #0001502).

## Acknowledgments

We are grateful for the support of the Albert Einstein College of Medicine Molecular Cytogenetics Core Facility, the Albert Einstein College of Medicine Genomics Core, and the Albert Einstein College of Medicine Flow Cytometry Core Facility, all of which are funded by a grant from the National Cancer Institute (2P30CA013330).

## Author Contributions

RO, KJ and DML conceived and designed experiments and analyses, wrote the draft and edited the final manuscript. MH performed in vitro experiments. JZ conceived and designed the cell cycle flow experiment. DP and PC (second to last author) performed RNA-seq data analysis. YT and TMS prepared samples and performed RNA-seq. PC (eighth author) performed *in vivo* experiments and analyses. JF supplied the DFMO. RAC provided scientific feedback and revised the manuscript. All authors edited the final manuscript.

## Data and materials availability

All data associated with this study are present in the paper or the Supplementary Materials.

